# Urine proteomic analysis of the rat startle model

**DOI:** 10.1101/2022.11.19.517176

**Authors:** Chenyang Zhao, Yuqing Liu, Youhe Gao

## Abstract

Urine proteomics was applied to explore whether the effect of startle can be detected in urine. A combination of natural enemy odor and sound stimulation was used to establish the rat startle model. Urine samples were collected before and after startle, and urine proteomes before-after startle of each rat were compared individually. Regulatory subunits of glutamate-cysteine ligase was identified as the sole differential protein among all five startled rats. To our surprise, its functional partner catalytic subunits of glutamate-cysteine ligase was also identified in four out of five rats as differential protein. When comparing before-after startle as two groups, 22 differential proteins were identified which represent biological pathways including neurotransmitter transport and glucose transmembrane transport.

## 1. Introduction

Startle is a behavior that often occurs in life. As the saying goes, “once bitten, twice shy”, different degrees of fright will have a certain impact on our lives. There are no physiological indicators related to fright, and no research has observed the changes caused by fright in urine. This study used urine proteomics to explore whether this startle can be detected in urine. In the study of urine proteomics, most of them have used urine proteomics methods to find disease-related biomarkers. Compared with blood, the most widely used marker source, urine is the place that most of the wastes in blood are dumped into^1^. Its content is not strictly regulated by homeostasis mechanisms in the body and thus it tolerates changes to a much higher degree^1^. The urine proteome is relatively less complex, and changes characteristic of low-abundance proteins are more easily detected^2^. Therefore, urine is a good source of biomarkers. Because the urine proteome is susceptible to a variety of factors, such as diet, drug therapy, and daily activities, to make the experimental results more accurate, the key is to use a simple and controllable system. Because the genetic and environmental factors of animal models can be artificially controlled to minimize unrelated influencing factors, the use of animal models is a very appropriate experimental method.

Many studies have shown that urine has great application in the search for biomarkers of psychiatric disorders, such as parkinsonism^3^, Alzheimer’s disease^4^, depression^5^ and other diseases. Therefore, we established a rat startle model using a combination of natural enemy odor and sound stimulation to establish a rat startle model. Urine samples were collected before and after startle, and proteomics analysis was performed to explore whether this startle mood could be detected in urine by urine proteomics.

## 2. Materials and Methods

### 2.1 Rat model establishment

Five male Sprague—Dawley (SD) rats (160±20 g) were purchased from Beijing Vital River Laboratory Animal Technology Co., Ltd. All animals were housed with free access to a standard laboratory diet and water with indoor temperature (22±1 °C) and humidity (65∼70%). All animal protocols performed during the experiments were approved by the Institutional Animal Care and Use Committee of Peking Union Medical College (Approval ID: ACUC-A02-2014-008). All methods in this research were performed in accordance with the guidelines and regulations.

The animal model of startle was established as follows: Each of the 5 rats was separately placed in fresh cat litter mixed bedding in a self-controlled way, and high-intensity sound stimulation was performed at the same time every day. The performance and behaviors of the rats were recorded.

### 2.2 Urine collection

Rats were placed in metabolic cages to collect urine for 12 h before startle treatment, and then the rats were returned to a standard environment for 3 days before performing the startle experiments. Once a day for 4 consecutive days, the rats were placed in metabolic cages to collect urine for 12 h with the startle treatment performed on the fourth day. Rats were fasted and deprived of water during urine collection. The urine samples were stored at -80 °C immediately after collection.

### 2.3 Sample preparation

Urine protein extraction and quantification: Rat urine collected at two time points was centrifuged at 12000 × g for 40 min at 4 °C, and the supernatant was transferred to a new EP tube. The supernatant volume was tripled and mixed evenly with precooled ethanol, and the mixture was precipitated overnight at -20 °C. The next day, the ethanol supernatant mixture was centrifuged at 12,000 × g for 30 min at 4 °C, the supernatant was discarded, the protein precipitate was left, inverted onto filter paper, and dried with a hair dryer. After that, the protein precipitate was resuspended in 120 μL lysate (8 mol/L urea, 2 mol/L thiourea, 25 mmol/L dithiothreitol, 50 mmol/L Tris), blown repeatedly through the tip of a pipette until no solid precipitate was found, and thoroughly mixed in a vortex mixer for 2 h. After mixing well, the mixture was centrifuged at 12,000 × g for 30 min at 4 °C, and the supernatant protein was placed in a new EP tube to measure the protein concentration by the Bradford assay. The protein samples were stored at -80 °C for later use.

Urine protease cutting: One hundred micrograms of urinary proteins from each sample was added to the filter membrane of a 10 kDa ultrafiltration tube (Pall, Port Washington, NY, USA), which was placed in a 1.5 mL centrifuge tube. Then, 25 mmol/L NH_4_HCO_3_ solution was added to a total volume of 200 μL, and 20 mM dithiothreitol solution (DTT, Sigma) was added and it was heated at 97 °C for 5 min and cooled naturally to room temperature. Then, 50 mM iodoacetamide (IAA, Sigma) was added and mixed at room temperature for 40 min in the dark. After the reaction finished, the proteins were processed in the following order: ① 200 μL of UA solution (8 mol/L urea, 0.1 mol/L Tris-HCL, pH 8.5) was added to the empty ultrafiltration tube and washed twice by centrifugation at 14000 × g for 5 min at 18 °C; ②Loading: protein samples were added to the treated ultrafiltration tube and centrifuged at 14000 × g for 40 min 18 °C; ③200 μL UA solution was added and centrifuged at 14000 × g for 40 min at 18 °C, repeated twice; ④ 25 mmol/L NH_4_HCO_3_ solution was added and centrifuged at 14000 × g for 40 min at 18 °C, repeated twice; ⑤Resuspended the denatured proteins with 25 mmol/L NH_4_HCO_3_ and digested with trypsin (Promega, Fitchburg, WI, enzyme to protein ratio of 1:50), 37 °C water bath overnight and finally passed through Oasis HLB cartridges (Waters, Milford, MA) for salt removal, placed in a SpeedVac (Thermo Fisher Scientific, Bremen, Germany) for drying, and then stored at -80 °C.

### 2.4 LC—MS/MS analysis

The digested samples were reconstituted with 0.1% formic acid, and the peptides were quantified using a BCA kit, diluting the peptide concentration to 0.5 μg/μL. Mixed peptide samples were prepared from 9 μL of each sample and separated using a high pH reversed-phase peptide separation kit (Thermo Fisher Scientific) according to the manufacturer’s instructions. Ten effluents (fractions) were collected by centrifugation, dried using a vacuum dryer and reconstituted with 0.1% formic acid in water. The iRT reagent (Biognosys, Switzerland) was added at a ratio of 1:10 v/v to all peptide samples to calibrate the retention time of the extracted peptide peaks. One microgram of each sample (a single experimental sample and ten fractions) was subjected to mass spectrometry and data acquisition using an EASY-nLC1200 Chromatography System (Thermo Fisher Scientific, USA) and Orbitrap Fusion Lumos Tribrid Mass Spectrometer (Thermo Fisher Scientific, USA).

### 2.5 Database searching and label-free quantitation

Ten raw files of the fractions were analyzed by PD (Proteome Discoverer 2.1) software, and the analysis results were used to establish the DIA acquisition method. The newly established DIA method is used to perform DIA mode acquisition for individual samples. Mass spectrometry data were processed and analyzed using Spectronaut X software at the end of acquisition. Raw files collected by DIA were imported for each sample for the search. Protein quantitation was performed using all fragment ion peak areas of the secondary peptides.

### 2.6 Statistical analysis

Differential proteins were screened by comparing the identified proteins before and after startle in a self-controlled manner. We screened the differential proteins under the following conditions: fold change between groups ≥ 1.5 or ≤ 0.67 and P value < 0.01 by paired t test analysis. The UniProt website was used for identifying the selected differential proteins (Fig. https://www.uniprot.org/), and the DAVID database (Fig. https://david.ncifcrf.gov/) was used to perform biological analysis. Functional analysis of the differentially expressed proteins was performed by searching the reported literature in the PubMed database.

## 3. Results and Discussion

### 3.1 Analysis of urine proteome changes before and after startle

### 3.1.1 Urine protein identification and unsupervised cluster analysis

After the rat startle model was established, a total of 10 urine protein samples collected before and after the startle were analyzed by liquid chromatography coupled with tandem mass spectrometry, and 1294 proteins were identified (≥ 2 specific peptides and FDR < 1% at the protein level). Total protein was analyzed by hierarchical clustering analysis (HCA), and the analysis results (Figure 2) showed that the group differentiation before and after the startle was very obvious.

**Figure 1.**
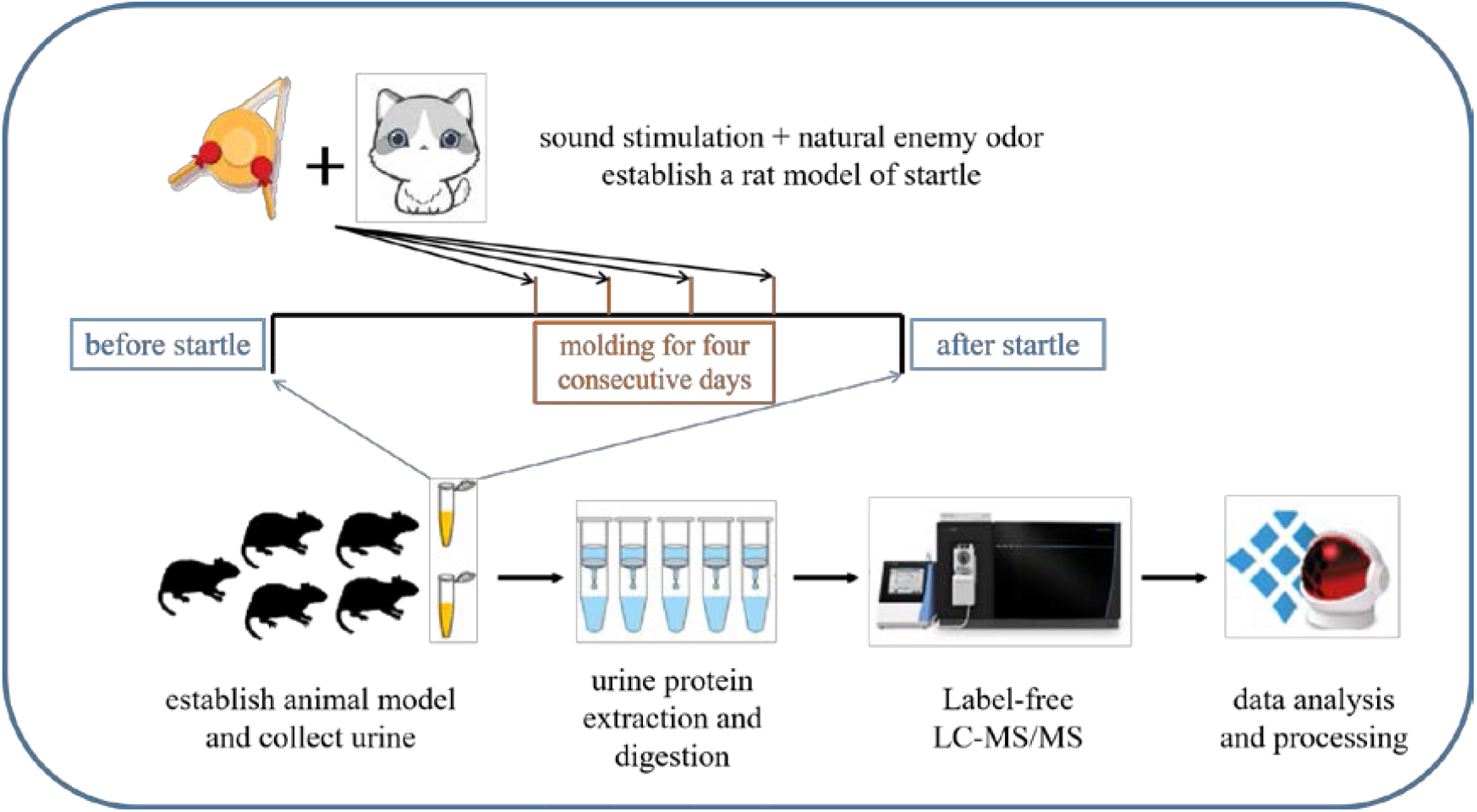
Technical flowchart of urinary protein identification of startled rats. Urine samples were collected, extracted, digested, and identified by liquid chromatography coupled with tandem mass spectrometry (LC—MS/MS) identification. Functional analysis of the differential proteins was performed by GO and IPA.

**Figure 2.**
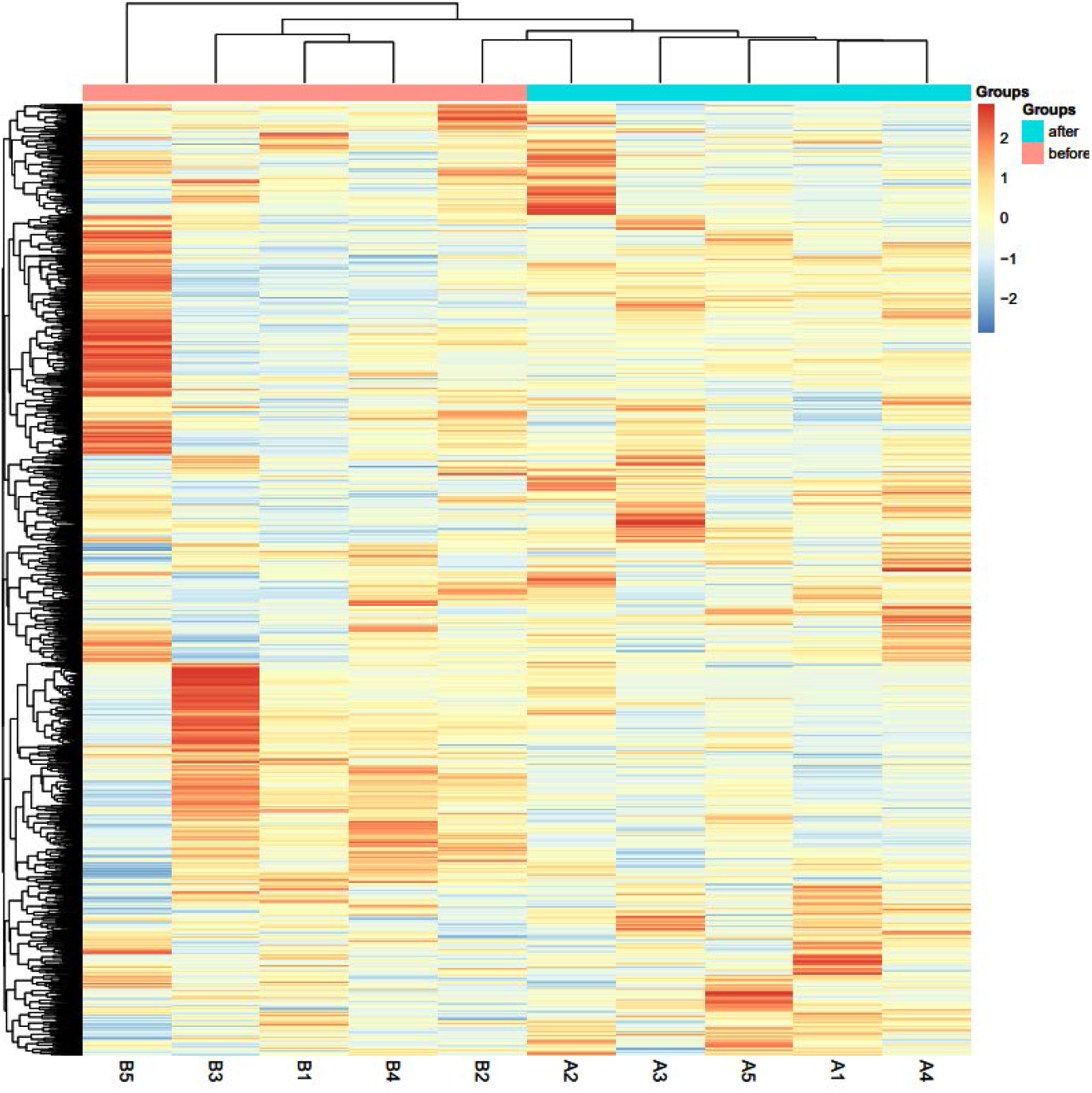
HCA analysis of identified urine proteins

**Figure 3.**
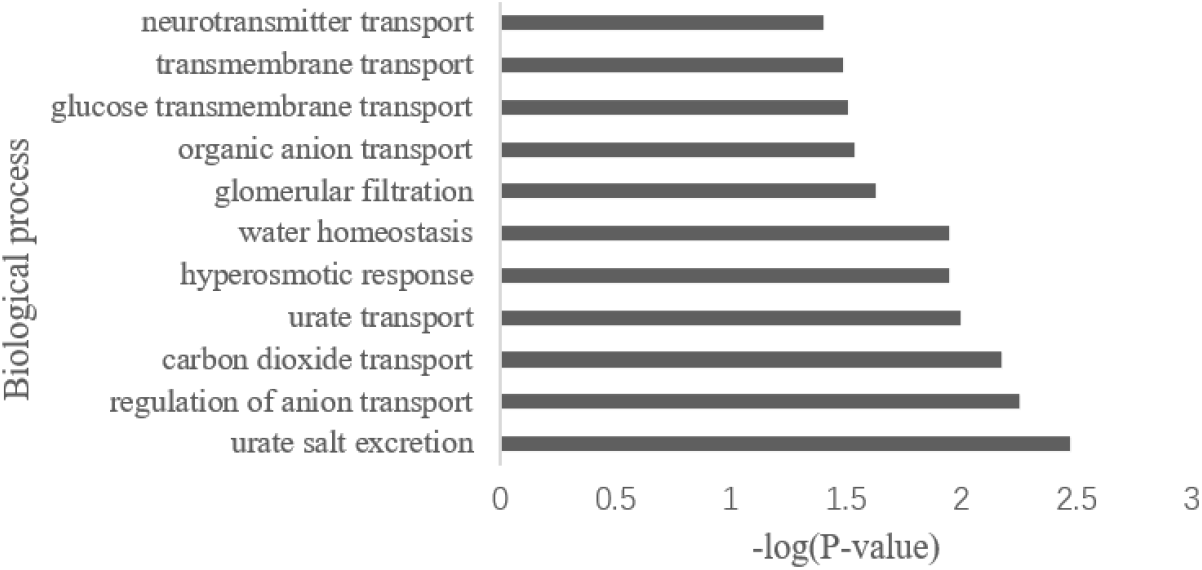
Biological process analysis of differentially expressed proteins

Urine protein before and after the startle was compared, and the criteria for screening the differential proteins were as follows: fold change ≥ 1.5 or ≤ 0.67 between groups, P < 0.01 by two-tailed unpaired t test. The results showed that 22 differential proteins could be identified after the startle compared with before the startle, and the specific information of the differential proteins is shown in Table 1.

**Table 1.** Details of the differentially expressed proteins

| Accession | Protein names | Fold change | Trend | P value |
| --- | --- | --- | --- | --- |
| F1LUS1 | Ig-like domain-containing protein | 7.58 | ↑ | 1.81E-03 |
| P61972 | Nuclear transport Factor 2 | 2.24 | ↑ | 1.40E-04 |
| G3V8Z5 | Ig-like domain-containing protein | 1.78 | ↑ | 9.83E-03 |
| D4A9Q5 | Carboxypeptidase M | 1.76 | ↑ | 6.28E-03 |
| Q6AXR4 | Beta-hexosaminidase subunit beta | 1.73 | ↑ | 4.21E-03 |
| P85971 | 6-phosphogluconolactonase | 1.62 | ↑ | 1.70E-03 |
| A0A0G2K3 W2 | Coagulation Factor V | 1.59 | ↑ | 9.50E-03 |
| G3V6V1 | Aminopeptidase B | 1.56 | ↑ | 1.79E-03 |
| Q7TPK2 | Ac2-120 | 1.55 | ↑ | 7.45E-03 |
| G3V647 | Pyridoxal kinase | 0.65 | ↓ | 5.73E-03 |
| P63029 | Translationally controlled tumor protein | 0.65 | ↓ | 7.57E-03 |
| P48500 | Triosephosphate isomerase | 0.63 | ↓ | 3.46E-03 |
| Q99MA2 | Xaa-Pro aminopeptidase 2 | 0.58 | ↓ | 5.82E-04 |
| Q5XI73 | Rho GDP-dissociation inhibitor 1 | 0.58 | ↓ | 6.86E-03 |
| P42854 | Regenerating islet-derived protein 3-gamma | 0.55 | ↓ | 9.73E-03 |
| P62749 | Hippocalcin-like protein 1 | 0.53 | ↓ | 6.00E-04 |
| Q9ESG3 | Collectrin | 0.49 | ↓ | 1.43E-03 |
| P46413 | Glutathione synthetase | 0.48 | ↓ | 6.50E-03 |
| P29975 | Aquaporin-1 | 0.46 | ↓ | 7.57E-04 |
| Q64319 | Neutral and basic amino acid transport protein rBAT | 0.33 | ↓ | 7.72E-03 |
| G3V8X5 | Solute carrier family 5 | 0.32 | ↓ | 5.49E-03 |
| A0A0G2JT43 | Solute carrier family 2 | 0.25 | ↓ | 4.98E-03 |

### 3.1.2 Different protein functional analyses

To investigate the function of these differentially expressed proteins, we performed functional analysis of the biological pathways using the DAVID database. The results showed that most of the pathways showed changes brought about by metabolism, such as urate metabolism and transport, CO_2_ transport, and organic anion transport. It is worth noting that these differentially expressed proteins were also enriched in biological pathways such as neurotransmitter transport and glucose transmembrane transport.

### 3.2 Analysis of urine proteome changes in individual rats before and after startle

To investigate whether the changes were consistent across the five rats, we performed urine proteomics analysis of self-controls before and after startle individually for each rat. Each rat’s urine profile was compared before and after startle. The conditions for screening the differential proteins were as follows: fold change ≥ 2 or ≤ 0.5, two-tailed unpaired t test P < 0.01. The results of the differential protein screening were as follows: 132 differential proteins were screened in rat 1, of which 44 proteins showed an upregulation trend and 88 proteins showed a downregulation trend; 79 differential proteins were screened in rat 2, of which 33 proteins showed an upregulation trend and 46 proteins showed a downregulation trend; 215 differential proteins were screened in rat 3, of which 38 proteins showed an upregulation trend and 177 proteins showed a downregulation trend; 91 differential proteins were screened in rat 4, of which 22 proteins showed an upregulation trend and 69 proteins showed a downregulation trend; and 134 differential proteins were screened in rat 5, of which 94 proteins showed an upregulation trend and 40 proteins showed a downregulation trend. We present a Wayne diagram of the differential proteins identified in the five rats (Figure 4), one of which was co-identified in all five, the glutamate-cysteine ligase regulatory subunit (P48508), and 19 proteins were co-identified in four rats. We listed the specific information and related studies of the human-derived proteins corresponding to these 20 proteins in Table 2. We found that these differentially expressed proteins were closely related to changes in nerve function, movement, metabolism, and blood pressure. In addition, glutamate-cysteine ligase regulatory and catalytic subunits were identified in all five or four of five rats, respectively, suggesting that these differential proteins are less likely to be produced by individual differences among the laboratory animals, and the differences we observed are more likely to be caused by fright.

**Figure 4.**
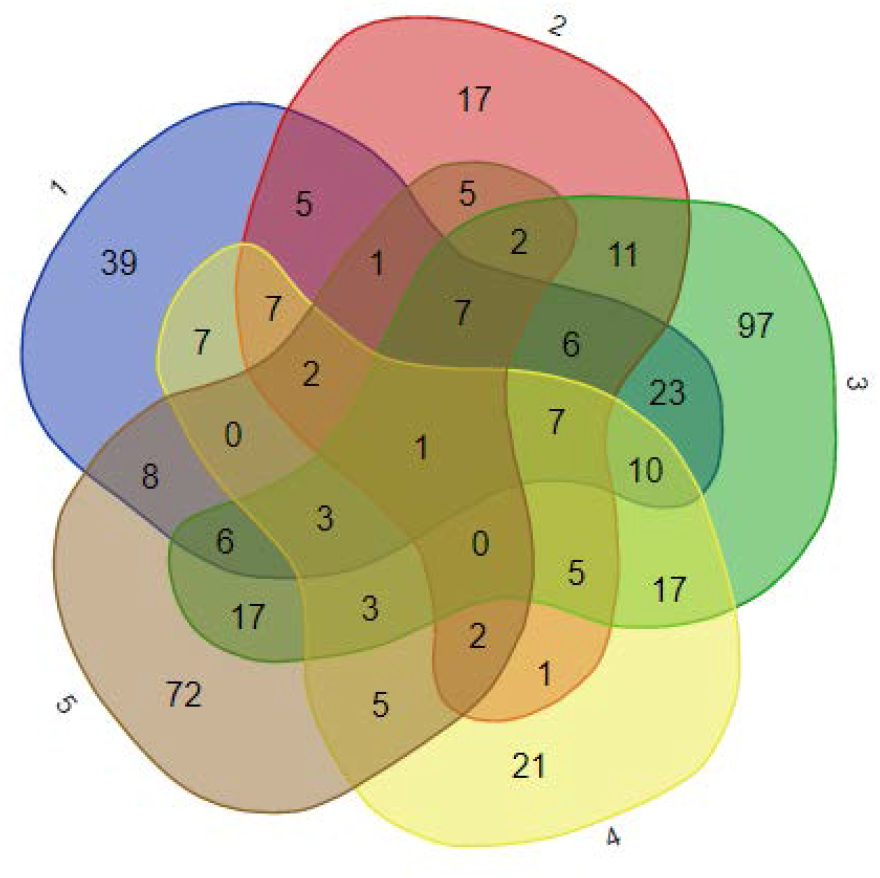
Venn diagram of differentially expressed proteins in a single rat

**Table 2.**
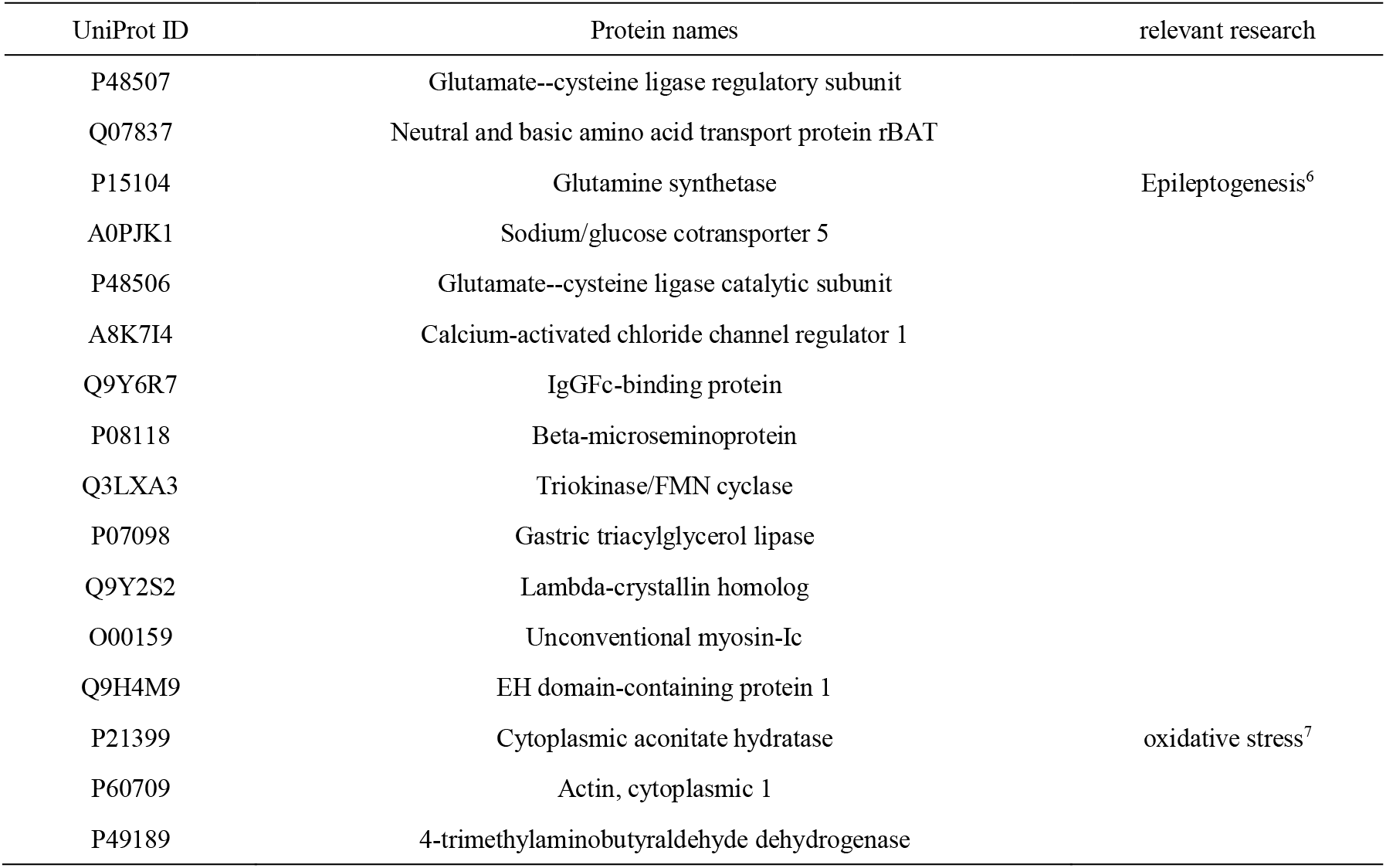
The co-identified differential proteins (*Homo sapiens*)

## 4. Conclusions

Regulatory and catalytic subunits of glutamate-cysteine ligase are both changed in urine after startle. Urinary proteins represent neurotransmitter transport and glucose transmembrane transport are also changed after startle. Urine proteome is sensitive enough to show effects of startle.

## CRediT authorship contribution statement

**Chenyang Zhao:** Conceptualization, Investigation, Writing-original draft, data curation, Visualization. **Yuqing Liu:** Investigation, Writing-original draft, Visualization. **Youhe Gao:** Conceptualization, Writing-review & editing, Supervision, Funding acquisition.

## Acknowledgements

This work was supported by the National Key Research and Development Program of China (2018YFC0910202); the Fundamental Research Funds for the Central Universities (2020KJZX002); the Beijing Natural Science Foundation (7172076)

